# Energy dissipation in early detection of cellular responses to metabolic challenges

**DOI:** 10.1101/2022.08.03.502646

**Authors:** Rafael N. Bento, Miguel A. Rendas, Valdir A. R. Semedo, Cátia F. Marques, Gonçalo. C. Justino, Carlos E. S. Bernardes, Manuel E. Minas da Piedade, Fernando Antunes

## Abstract

Metabolic alterations have been recognized to underly the etiology of many diseases. Herein, cellular energy dissipation was evaluated as a novel non-specific global biomarker of metabolic alterations. Energy dissipation, measured as heat by microcalorimetry, was maximal during *Saccharomyces cerevisiae* adaptation to growth conditions before fast proliferation took place. This response was further augmented by 95 % in media where nutrient assimilation was more difficult, and by 133 % under sub-optimal non-carbon nutrient levels. In this last case, the increase in energy dissipation (1) reflected changes in amino acid and glycolytic metabolism and (2) anticipated changes in the growth curve significantly later observed by traditional microbiological measurements. It was, therefore, an early marker of adaptive responses that compensated for sub-optimal nutrient levels and maintained phenotypic stability. Compensatory responses buffer systems against perturbations and delay the onset of diseases. Microcalorimetry can, therefore, provide a biomarker development platform for early disease-diagnosis.

**HIGHLIGHTS:**

1. Energy dissipation measurements detect cell responses to metabolic challenges.
2. The detection by microcalorimetry occurs considerably earlier than by traditional microbiological measurements.
3. Sub-optimal non-carbon nutrient levels impact energy dissipation long before cell proliferation.
4. Energy dissipation is highly sensitive to increased nutrient assimilation difficulty.

## INTRODUCTION

Metabolic alterations are at the core of many disease processes (DeBerardinis and Thompson, 2012). Currently, most disease biomarkers are specific for individual pathologies, being based on the detection of particular molecules (Wishart et al., 2021). Here, we present an alternative approach and propose energy dissipation (ED) as a global non-specific biomarker for early detection of metabolic alterations. Living systems respond to perturbations/stress, such as genetic or environmental alterations, by triggering adaptive mechanisms that include compensatory or redundant pathways to maintain phenotypic stability. Eventually, when subjected to long-lasting or high-intensity perturbations, cells are not able to maintain phenotype stability and diseases emerge (Hartman et al., 2001a). In general, response to perturbations is expected to have a significant energy cost, reflected by a highly dissipative character, because it entails a rapid change of the cellular state mediated by far from equilibrium processes (Lan et al., 2012; Woronoff et al., 2019). This can be rationalized by noting that according to thermodynamics the efficiency of chemical reactions within the cell would be maximal if they occurred reversibly or, in other words, through a sequence of quasi-equilibrium states separated by infinitesimal differences. Reversible processes correspond, however, to limiting cases, which do not occur in practice, since by requiring an infinite number of steps they are also infinitely slow. The larger the rate of response, the larger the departure from reversibility and the larger energy dissipation cost (Beard and Qian, 2007; Schaarschmidt et al., 1975). Thus, we hypothesized that ED could provide a sensitive indicator of responses to metabolic perturbations or challenges.

The model used to test this hypothesis was *Saccharomyces cerevisiae* (*S. cerevisiae*). *S. cerevisiae* is particularly appropriate because of its genetic redundancy, with only ~18% of genes being essential for growth on a rich medium (Giaever et al, 2002), and extensive functional redundancy in pathways (Harrison et al., 2007; Papp, 2007; Ulitsky and Shamir, 2007). These features give *S. cerevisiae* the ability to overcome several metabolic challenges by recurring to alternative metabolic pathways. We employed the common *S. cerevisiae* laboratory strain BY4741, which is auxotrophic for L-histidine, L-leucine, L-methionine, and uracil, due to deletion of the corresponding anabolic genes (Brachmann et al., 1998). Quiescent BY4741 cells were subjected to three metabolic challenges: (1) exposure to growth media to trigger high proliferative rates; (2) exposure to complex media to difficult nutrient assimilation; and (3) exposure to sub-optimal non-carbon nutrients. Cells were separately grown in Synthetic Complete (SC), enriched Synthetic Complete (eSC), and Yeast Peptone Dextrose (YPD) media. All three media contained non-limiting and identical levels of glucose (the main carbon source), but different concentrations of other nutrients needed for growth, including the auxotrophic nutrients. The complex YPD and the chemically defined SC media were originally developed to allow optimal growth of *S. cerevisiae* wild-type strains. It was, however, later concluded that in the case of BY4741 only sub-optimal growth conditions were provided by the SC formulation (Cohen and Engelberg, 2007; Corbacho et al., 2011; Hanscho et al., 2012). Thus, in eSC all nutrients, except glucose, had their concentrations doubled relative to the default SC medium. Finally, ED was measured by flow-microcalorimetry, a non-invasive and extremely sensitive technique (Beezer, 1980; James, 1987) that provides real time information on metabolic activity, through the heat released by organisms or cells (Beezer, 1980; James, 1987). In parallel, a metabolomic analysis was performed to complement the integrative ED data with molecular level information.

ED proved to be extremely sensitive to the three metabolic challenges imposed. For the challenge imposed by sub-optimal non-carbon nutrients, a large ED increment was observed long before traditional microbiological measurements were impacted. In conclusion, ED is a potential biomarker that signals response mechanisms to metabolic perturbations and can detect these alterations before phenotypic stability is lost.

## RESULTS

### ED microcalorimetric traces from whole cell cultures hide the impact of metabolic challenges during adaptation to growth conditions

To analyze the impact of metabolic challenges on the adaptation to growth conditions, samples of *S. cerevisiae* cultures in stationary phase (in which nutrients are fully exhausted) were transferred to fresh YPD, SC, or eSC media. In the SC medium, nutrient exhaustion occurs faster than in either YPD or eSC. Thus, according to our hypothesis, the first signs of compensatory/redundant pathways activation, reflected by a higher ED, should be observed earlier in the SC growth curve, namely, during the lag phase. The energy dissipation by *S. cerevisiae*, was experimentally determined as a power by flow-microcalorimetry. This power corresponds to a ED rate (*r*_ED_) and has units of W or J s^-1^. It is the calorimetric parameter traditionally used to follow the overall metabolic activity in cell cultures based on heat release measurements. In parallel, microbiological growth parameters, such as the duration of the lag phase and the maximum growth rate, were determined by optical density measurements at 600 nm (*OD*_600_). The *r*_ED_ calorimetric profile (Figure 1A) was similar to previously reported ones, with a well-defined lag phase, where power dissipation by the cellular culture was low, followed by the exponential phase where *r*_ED_ rapidly increased (Blomberg et al., 1988; Brettel et al., 1980). No noticeable differences were observed during the lag phase among the profiles acquired in the three distinct growth media. Only later during the growth cycle, differences were observed that clearly reflected media dependent metabolic changes. There was a significantly higher *r*_ED_peak maximum for YPD than for eSC and SC media. This observation agreed with the *OD_600_* reading (Figure 1B) when glucose was exhausted, indicating a higher yield of biomass per mol of glucose consumed in the YPD medium. As mentioned above, the BY4741 *S. cerevisiae* strain is auxotrophic for L-histidine, L-leucine, L-methionine, and uracil (Brachmann et al., 1998). These auxotrophies are associated with the deletion of the anabolic genes *HIS3*, *LEU2*, *MET15*, and *URA3*, and they were shown to originate media dependent growth patterns (Cohen and Engelberg, 2007; Corbacho et al., 2011; Hanscho et al., 2012). Because BY4741 cannot endogenously synthesize all the nutrients needed, differences in their initial concentration and rate of depletion from the growth medium originate a lower efficiency of glucose assimilation along the cell growth cycle. Growth in the default SC medium is, therefore, characterized by a briefer segment of the exponential phase where cells can proliferate at maximum division rate, reaching lower *r*_ED_peak maximum and final biomass produced.

**Figure 1.**
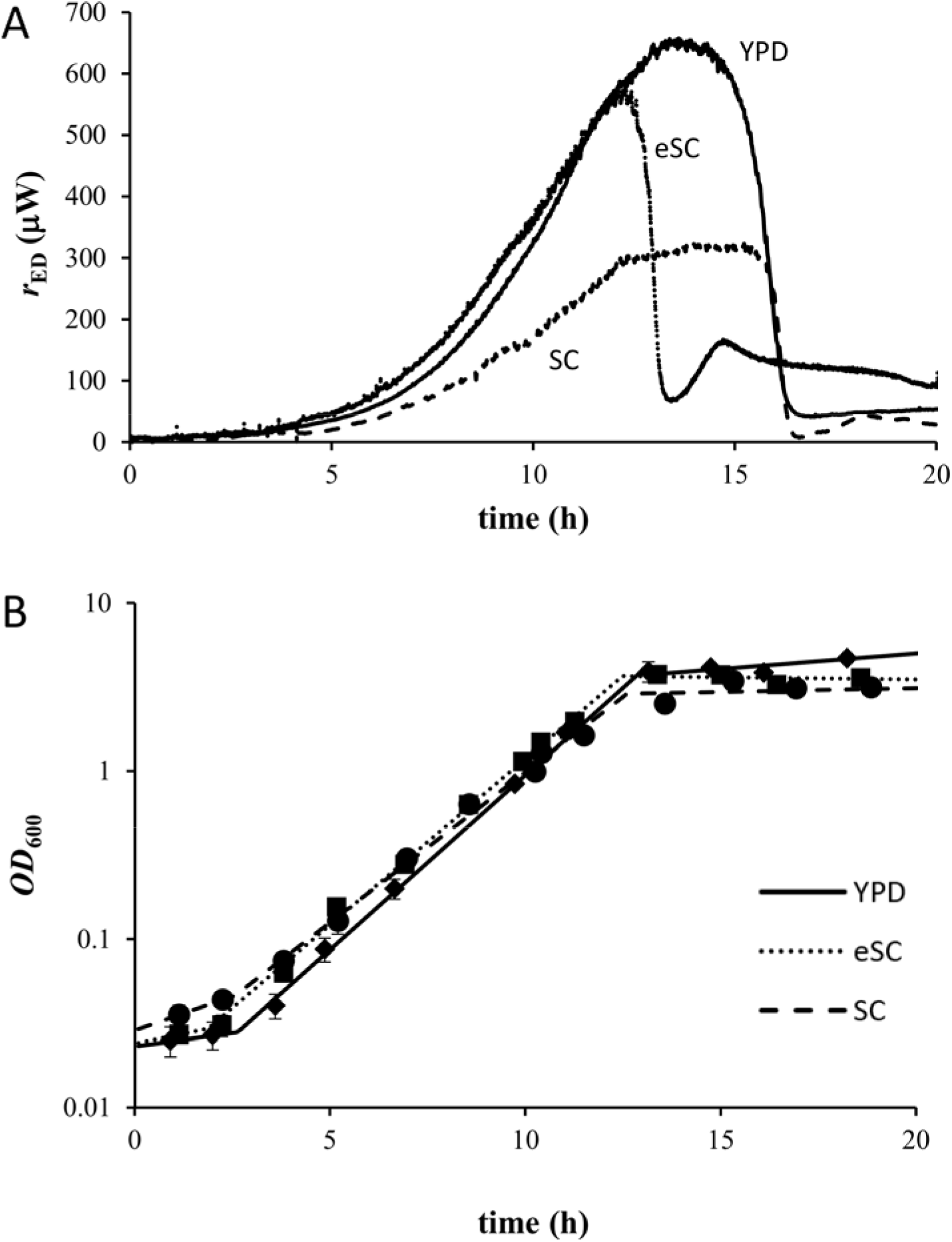
ED rates of the whole cell culture do not anticipate the impact of sub-optimal nutrient concentrations on adaptation to growth conditions. (A-B) The growth of *S. cerevisiae* cells in YPD (—), SC (- - -), and eSC (…..) media was followed by ED microcalorimetry measurements and by optical density measurements at 600 nm (*OD*_600_). Whole cell culture ED rates correspond to representative experiments. *OD*_600_ data points represent optical densities obtained at specific times (*n*= 5 for YPD, *n* = 5 for SC and *n* = 5 for eSC), and lines correspond to linear least squares fittings to those data.

In conclusion, signatures of media-dependent metabolic changes are noted in the *r*_ED_ curves. These observations cannot, however, be taken as a suitable earlier indicator of metabolic perturbations since these are only detected several hours after inoculation and are also concomitantly evidenced by microbiological measurements. The *r*_ED_ curve hides small changes in the early phase of the growth curve when a low number of cells is present in the culture. Analyses of growth curves based on ED rates alone can, therefore, be misleading as, for example, they apparently suggest that the lag phase corresponds to an intrinsic dormant state, where very little metabolic activity is going on compared to the exponential growth stage. It has, however, been recognized that, during the lag phase, microorganisms adapt to a new medium and, consequently, some significant metabolic activity is expected to take place (Schiraldi, 1995).

### ED rates normalized by the cell number are extremely sensitive to metabolic challenges and anticipate the impact of sub-optimal nutrient concentrations on cell growth

To uncover hidden information in the ED rate profile, we normalized the *r*_ED_ rates by the cell number, obtaining the specific ED rate, _sp_*r*_ED_, a parameter with units of W cell^-1^. _sp_*r*_ED_ profiles considerably differed from the corresponding *r*_ED_ traces and gave important insights into the metabolic dynamics along the *S. cerevisiae* growth cycle. As shown in Figure 2A, the _sp_*r*_ED_increased very rapidly during the lag phase. This could be expected because, when inoculated, cells are in a stationary quiescent phase, which has an associated low metabolic activity, resulting in a lower ED. Cells subsequently build-up the necessary machinery for growth and proliferation. This corresponds to a highly dissipative process, as reflected by the fast _sp_*r*_ED_increase noted in Figure 2A. One key aspect in Figure 2A is that, in agreement with previous observations (Robador et al., 2018; Schaarschmidt et al., 1975), the maximum value of the _sp_*r*_ED_in all growth media corresponds to the transition from the lag to the exponential phase. In contrast with what could be inferred from the *r*_ED_ of whole cell cultures, at the cellular level, the lag phase displays the most intense dissipative metabolism within the growth cycle. Quantitatively, the maximal _sp_*r*_ED_ during the lag phase was 2 to 4 times higher than that observed at the end of exponential phase, when active cell proliferation started to decrease from its maximum value (see Figure 2B). This may be caused by the break of detailed balances during feed-back loops associated with adaptation to a new medium, which entails large deviations of metabolism operation from reversibility and hence high energy dissipation (Lan et al., 2012). Thus, adaptation to the growth conditions in the lag phase is highlighted by a significant increase in ED, even when compared with the ED observed for maximal proliferation rates during the exponential phase.

**Figure 2.**
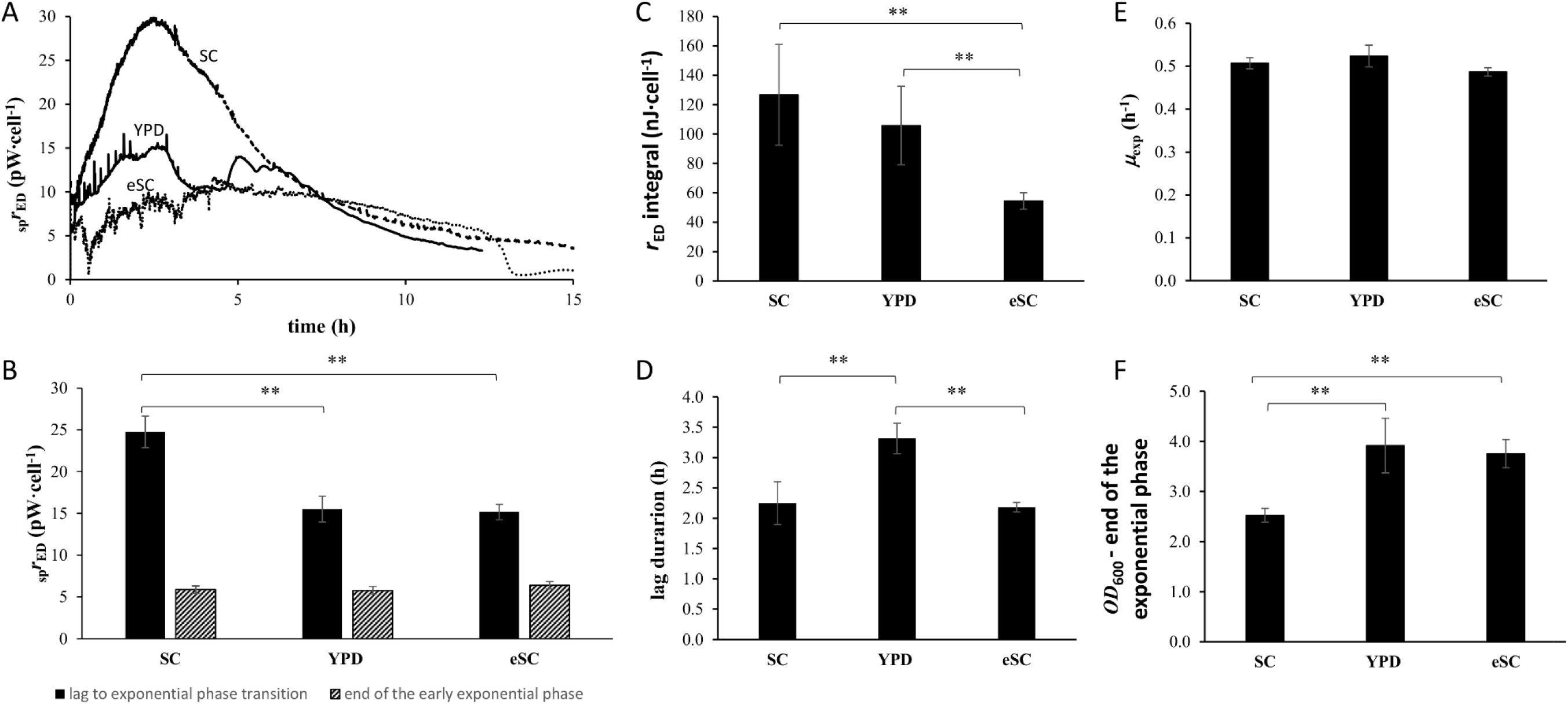
ED rates normalized by the cell number are extremely sensitive to metabolic challenges and anticipate the impact of sub-optimal nutrient concentrations on cell growth. (A) The specific ED rate for cultures grown in YPD (—), SC (- - -), and eSC (…..) media. Data corresponds to representative experiments. (B) The specific ED rate obtained during the transition from the lag to the exponential phase and at the end of the early exponential phase are compared for YPD (*n*=7-10), SC (*n*=8), and eSC media (*n*=3-5). (C) Integral of *r*_ED_ along the lag phase normalized by the number of cells. (D) The length of the lag phase in YPD (*n*=7), eSC (*n*=5), and SC media (*n*=7). (E) The maximum proliferative rate in YPD (*n*=9), eSC (*n*=9), and SC media (*n*=10). (F) The *OD*_600_ value at the end of the exponential phase in YPD (*n*=5), eSC (*n*=4), and SC media (*n*=5). In Figures (B-F), data are represented as mean ± SD of the number of independent determinations. ** *p* < 0.01 and * *p* < 0.05

Can the challenge imposed by the complexity of the YPD media be detected by ED alterations? YPD media includes yeast extract, and is, therefore, a more complex medium than the simpler and well-defined eSC medium. In YPD additional molecular machinery must be produced to assimilate the more complex nutrients present (Melnykov, 2016; Proust et al., 2019). This could, in principle, result in higher ED during the lag phase. While the maximal _sp_*r*_ED_ at the transition from the lag to the exponential phase was similar in YPD and eSC (Figure 2B), the overall energy dissipation during the lag phase, calculated as *r*_ED_ integral along the whole lag phase normalized by the number of cells, was 94 % higher in YPD (105.78 ± 26.69 nJ cell^-1^, *n* = 5) than in eSC (54.44 ± 5.63 nJ cell^-1^, *n* = 4) (Figure 2C). The higher ED in the lag phase reflects the longer duration of this period for cells grown in YPD (3.31 ± 0.25 h, *n* = 7) than for those grown in eSC (2.18±0.08 h, *n* = 5) (Figure 2D). The observation of a longer lag phase in YPD probably resulted from the buildup of additional molecular machinery needed to assimilate nutrients from the more complex YPD medium. Thus, even if the maximal _sp_*r*_ED_ are similar in YPD and eSC, higher energy dissipation along the lag phase is required for adaptation to the former than to the latter medium. Despite these differences, the similar maximal proliferation rates observed for YPD (0.52±0.03 h^-1^, *n* = 9) and eSC (0.49±0.01 h^-1^, *n* = 9) (Figure 2E) gave a clear indication that cells adapted equally well to both media. Thus, specific ED measurements detected the cellular response triggered by an increased difficulty in assimilating nutrients from the environment.

ED was impacted by (1) the increased cellular activity associated with adaptive processes in the lag phase and by (2) the harder challenge imposed to cells by complex media during adaptation to growth conditions. Can ED also detect the increased adaptive effort caused by sub-optimal nutrient levels in the SC medium? To answer this question, ED was compared in the eSC and SC media only, since the complexity associated with the presence of yeast extract in YPD media would introduce a variability that is difficult to characterize. The very high level of the _sp_*r*_ED_ observed at the transition from the lag to the exponential phase was 63% higher in SC (24.76±1.88 pW cell^-1^, *n* = 8) than in eSC (15.17±0.93 pW cell^-1^, *n* = 3) media (Figure 2B). In addition, the total energy dissipated during the lag phase calculated by integrating _sp_*r*_ED_ along the whole lag phase was 133% higher in SC (126.70±34.24 nJ cell^-1^, *n* = 5) than in eSC (54.44±5.63 nJ cell^-1^, *n* = 4) (Figure 2C). Thus, a sub-optimal nutrient content triggered an early cellular response that was detected as a large ED increase during the lag phase, where the overall depletion of nutrients should be negligible.

Do changes in ED during the lag phase anticipate alterations in traditional microbiological growth parameters? As referred to above, at the end of the exponential phase the *OD_600_* reading was higher in the richer eSC than in the poorer SC media. This indicated a higher yield of biomass per mol of glucose consumed in the former medium (Figure 2F). Despite this observation, similar maximum growth rates (*μ*_exp_) driven by a respiro-fermentative metabolism were observed in SC and eSC after the lag phase was concluded (Figure 2E). The obtained proliferation rates were 0.51±0.01 h^-1^ (*n* = 10) for SC and 0.49±0.01 h^-1^ (*n* = 9) for eSC, indicating that cells entered the exponential phase equally adapted to growth conditions in both media. In addition, the duration of the lag phase was also similar in SC (2.25±0.35 h, *n* = 7) and eSC (2.18±0.08 h, *n* = 5) (Figure 2D). Therefore, the impact of sub-optimal nutrient conditions upon traditional microbiological parameters was visible only at latter stages of the growth curve as a shortening of the exponential phase. In conclusion, while specific ED changes already signaled the effect of sub-optimal nutrient levels during the lag phase, traditional microbiological measurements would only detect the effect later through a shortening of the exponential phase. entailing

### Sub-optimal nutrient concentrations decrease intracellular amino acid and glycolytic pools

The changes in ED during the lag phase described in the previous section sum up each metabolic change implicated in the response to nutrient limitations. To elucidate the molecular changes behind the observed increase in energy dissipation, a metabolomic analysis during the lag phase and early part of the exponential phase was performed. This analysis was restricted to cells growing in eSC and SC media, because the complexity of the YPD components would introduce confounding variables.

We started by analyzing amino acid levels, since their biosynthesis has a high energy demand. Because SC media contains half of the amino acid content of eSC media, we evaluated if this difference was reflected in the intracellular pool. The two media showed no significant statistical difference in the intracellular amino acid pool at 1 h (Fig. 3A). For longer time courses, however, amino acid levels were decreased by 46 % at 2 h (p < 0.05) and 81 % at 5 h (p < 0.01) in the SC exhibited (Fig. 3A). Next, we performed an analysis of the dynamics of each amino acid during the first 5 hours of the growth cycle (Fig. 3B and C). It was observed that: (*i*) the levels of Gly and Lys were very resilient in both media throughout this period; (*ii*) the Asp, Ser, Met, Ala, and Pro levels were essentially identical to those observed at 1 h in eSC medium but decreased at 2 h and/or 5 h in the poorer SC medium; (*iii*) finally, the Val, Ile, Tyr and His levels were found to be reduced at 5 h in the eSC medium, while in the poorer SC medium the decrease was already felt at 2 h, and further intensified at 5 h. In summary, during the first 5 h of growth, the decay of the amino acid intracellular pool was much more pronounced in the SC than in eSC medium.

**Figure 3.**
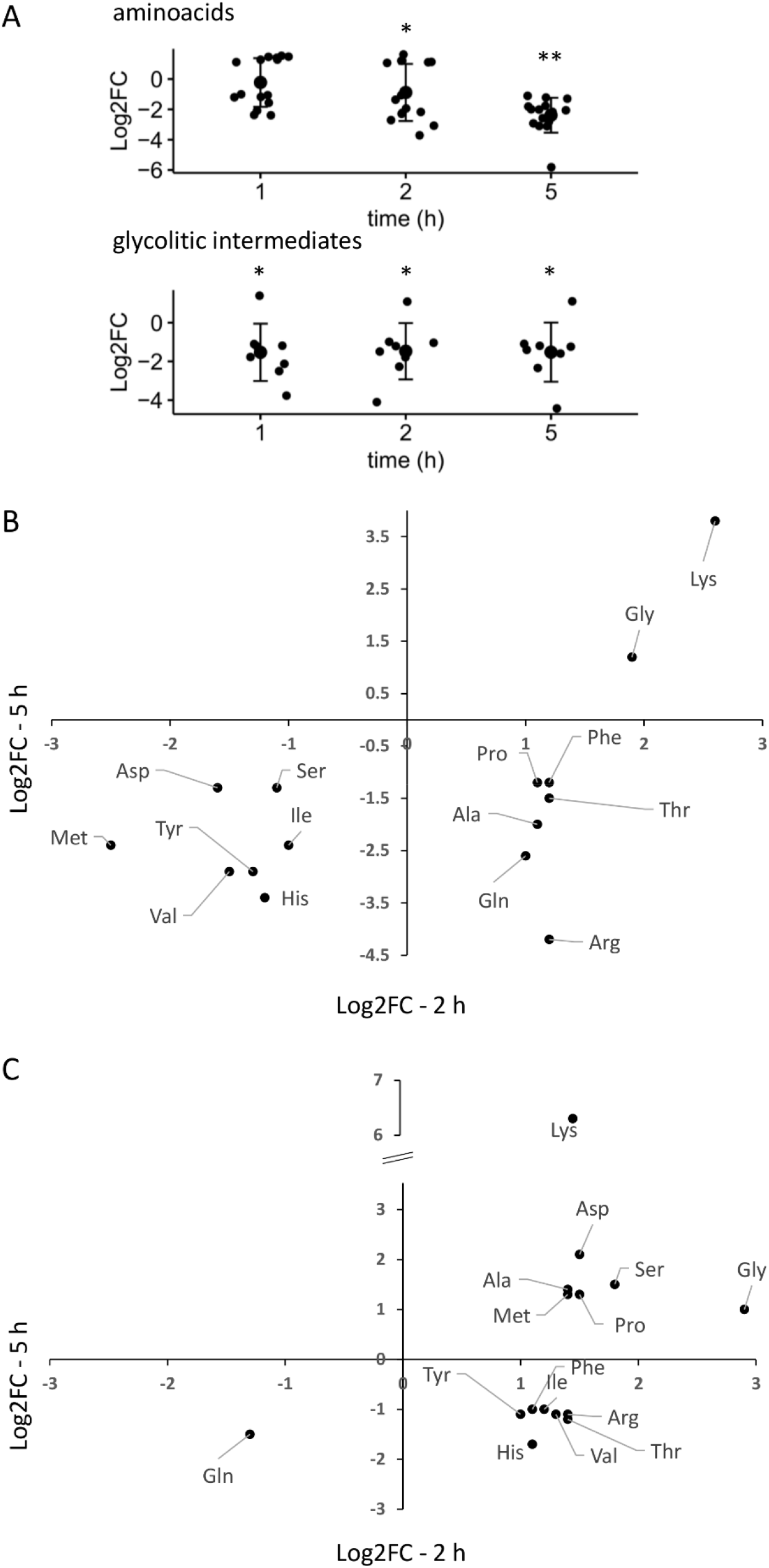
Sub-optimal nutrient concentrations decrease intracellular amino acid and glycolytic pools. (A) Log2 fold-changes of intracellular metabolite levels for cultures grown in Sc compared with eSC media during the lag phase (1 h), at the transition from the lag to the exponential phase (2 h), and at the exponential phase (5 h). Individual metabolites alterations can be seen in SI (Figure SI1). \*\**p* < 0.01 and \**p* < 0.05 (B,C) Dynamics of individual amino acids in SC and eSC media. Log2 fold-changes at 5 h (y-axis) and 2 h (x-axis), both referenced to metabolite levels at 1 h, are plotted. Only metabolites with a significant change are shown. Log2 fold-change and significance cutoffs for differential levels were 1 (or −1) and 0.05, respectively. Fold-changes are the average of three independent experiments.

Concerning the glycolytic pathway, albeit the initial glucose availability to *S. cerevisiae* cells was the same in both media, the set of all glycolytic metabolites identified was already significantly depressed in SC compared to eSC medium at 1 h. This tendency persisted throughout the period analyzed (Fig. 3A). Of relevance, were also the large decreases observed in hexoses (>75 % decrease) and fructose-1,6-biphosphate (>90 % decrease) levels. This indicates that the glycolytic metabolism is altered since the very beginning of the growth cycle, even if the initial carbon source availability is similar in both media. It should also be noted that no clear difference in the levels of citric acid cycle intermediates was observed between the two media (SI), probably reflecting the amphibolic role of this cycle.

In conclusion, metabolic and ED variations were concomitantly identified much earlier in the growth cycle than microbiological changes.

## DISCUSSION

The central idea that biological systems increase ED when adapting to alterations of the conditions in which they operate, directly results from thermodynamic laws. But are these increases measurable against background metabolic activity? Can they be used to detect biochemical deficiencies earlier? This study shows for the first time that ED is an extremely sensitive parameter for unveiling cellular responses to metabolic challenges. We demonstrated that _sp_*r*_ED_ increased significantly during adaptive response to growth conditions, being 2 to 4 times higher at the end of the lag phase than at the end of the high proliferative exponential phase. In addition, higher complexity of growth media increased the overall ED in the lag phase by 94%, impacting cellular function and causing a loss of phenotypic stability manifested by a longer period for cells to adapt to growth conditions. Lastly, sub-optimal non-carbon nutrient levels increased the overall ED in the lag phase by 133%, and the maximal *r*_ED_ by 63%, concomitantly with diminished amino acid and glycolytic intracellular pools. Phenotypic stability was, however, maintained for a longer period, as the duration of the cellular adaptation to growth conditions and the maximal rate of cell proliferation were not influenced. The observed ED increase, long before phenotypic stability was impacted, holds a high potential for the future use of ED as a sensitive early biomarker of defects that are initially phenotypic silent. This characteristic is highly relevant because buffering of challenges is a common biological response. In large-scale genetic studies, titration of gene expression shows that the capacity to buffer perturbations up to a certain threshold is very common (Jost et al., 2020; Keren et al., 2016). At the systems level, perturbations through the action of the cellular network are expected to be even higher than those found for individual genetic perturbations (Hartman et al., 2001b). In degenerative diseases, synaptic changes are buffered to keep functionality long-before the onset of disease (Orr et al., 2020). In the present work, amino acid and glycolytic pool decreases were buffered for several hours before cellular growth was affected. Thus, ED is a potential biomarker of cell functionality, with cells dissipating more energy to perform a certain function when they are less fit. Eventually these cells become dysfunctional, and the pathological process accelerates. Anticipating the detection of functional alterations by an ED increase holds a high potential for biomarker development. But how general are the responses observed in this work? A key characteristic of this work is the presence of abundant levels of carbon energy sources that were similar in all media. This energetic abundancy probably enabled the increased ED in response to a challenge, to maintain cellular performance. Previously, it was reported that during adaptation to a chemical stress under conditions of progressive energy and carbon source scarcity, cells maintain an adaptive response but take longer to achieve adaptation (Lan et al., 2012). In face of challenges, two possible strategies for resource allocation are: (1) sparing resources albeit at a cost of performance, as observed in (Lan et al., 2012); or (2) maintain function at a cost of faster ED as observed here. This last strategy is a good example of phenotypic stability that is probably enabled by the abundant carbon energy sources employed in this work. This is relevant for the application of ED as an early disease biomarker in the modern environment of constant high resource availability.

### Final remarks

We propose that ED constitutes a biomarker of cellular responses to genetic/environmental perturbations, fulfilling a potential role as universal biomarker of the cellular effort to fulfill their functions. A higher ED indicates that the cell is less fit to fulfill its function, and thus dissipates more energy. Eventually, this lower fitness will result in a loss of function and the emergence of pathology. Thus, pathologies that impact different cell types could be early detected by evaluation the ED rate in those cells. In an era of big data where large information sets are obtained by omic approaches, a single measure of the global state of the cell can work as a simple and effective surrogate endpoint in early disease diagnostic.

## ACKNOWLEDGMENTS

This work was supported by Fundação para a Ciência e a Tecnologia (FCT), Portugal through PIDDAC (UIDB/00100/2020 and UIDP/00100/2020), LA/P/0056/2020), RNEM-LISBOA-01-0145-FEDER-022125, SFRH/BD/117787/2016 (PhD fellowship to RNB) and 2021.03239.CEECIND (CEEC fellowship to CESB). We thank Dr. Hermínio P. Diogo for his contribution to the design of the cell culture vessel adapted to the calorimetric apparatus.

## AUTHOR CONTRIBUTIONS

RNB, MEMP and FA designed the experiments and wrote the manuscript with input from all authors. RNB perform all calorimetry experiments except as noted. VARS and MAR optimized the calorimetric set up. CFM and GCJ performed all metabolomic experiments. CESB developed methods and in house instrumentation for calorimetric experiments.

## DECLARATION OF INTERESTS

The authors declare no conflicts of interest.

## METHODS

### Yeast strain, media, and growth conditions

*Materials. S. cerevisiae* haploid strain Y00000 (wild-type, genotype BY4741 *MATa his3Δ1 leu2Δ0 met15Δ0 ura3Δ0*) (EUROSCARF, Frankfurt, Germany). Yeast extract, bactopeptone, yeast nitrogen base and agar (Becton Dickinson and Company, Le Pont de Claix, France), glucose (Merck, Darmstadt, Germany), amino acids and nitrogen bases (Sigma Chemical Company, St. Louis, MO, USA), and water from a Milli-Q^®^ Type 1 Ultrapure Water system.

*Media and growth conditions*. Growth and all incubations were carried out in either (*i*) Yeast Peptone Dextrose (YPD) containing 0.9% (w/v) yeast extract, 1.8% (w/v) bactopeptone and 2% (w/v) D-glucose; (*ii*) in Synthetic Complete (SC) medium containing 0.685% (w/v) yeast nitrogen base, 2% (w/v) D-glucose, and the amino acids and nitrogen bases as indicated in (Branco et al., 2004); or. (*iii*) enriched Synthetic Complete (eSC) medium containing 2% (w/v) D-glucose and doubled concentrations of all other components compared with SC medium.

Cells were growth at 30 °C under aerobic and batch in an Infors AG CH-4103 Bottmingen incubator under shaking at 160 rpm. Experiments were initiated by inoculating in growth medium stationary phase *S. cerevisiae* cells at an initial *OD*_600_ in the range 0.02-0.03. Growth was followed by optical density at 600 nm (OD_600_) and by counting cells in a hemocytometer. A value of 1 OD_600_ = (5.70±0.11)×10^7^ cells·mL^-1^ was obtained.

### Calorimetric system

Calorimetric experiments were carried out with a modified version of the previously described LKB 10700-1 flow calorimeter operating in the flow-through mode (Leskiv et al., 2009). The modifications were essentially related to the adaptation of ancillary equipment for cell growth and circulation though the calorimetric sensor. Cell cultures were kept in a jacketed glass vessel that was placed inside an incubator adjacent to the calorimeter. Appropriate oxygenation was ensured by magnetic stirring. Constant humidity was achieved by placing an open vessel filled with distilled water inside the incubator. The cell cultures were circulated between the glass vessel and the calorimetric cell through Teflon tubes. Pumping was by means of an Ismatec MS-4/12 multi-channel peristaltic pump located inside the incubator. The whole system was thermostated at 30 °C, including the Teflon tubes used to transport the cell culture between the glass vessel and the calorimetric cell. This last feature was achieved by keeping the Teflon lines exiting the incubator and returning from the calorimeter inside flexible PVC pipe bellows, where thermostated air from the incubator was circulated by a system of fans. Different thermostatic units ensured a temperature stability of ±0.01 K in the glass vessel (HAAKE DC 5 unit), ±0.03 K in the incubator (Julabo LC6 controller), and ±0.008 K in the calorimeter (LKB 10700-1 air thermostat jacket). The thermostatization of the glass vessel inside an already temperature-controlled environment allowed a better stabilization of the cell culture temperature, which became essentially unaffected by the small variations associated with the opening/closing of the incubator during the experiments. The calorimetric apparatus was kept in an airconditioned room whose temperature was regulated to 22±1 °C. The volume of culture in the reaction vessel was 20 mL. The nominal volume flow rate in the peristaltic pump was adjusted so achieve an actual flow rate of 1.0 mL·min^-1^. This adjustment was regularly checked, since flow rate variations affect various calorimetric parameters, such as the calibration constant (Leskiv et al., 2009), the effective volume of the calorimetric vessel (Bento et al., 2018; Leskiv et al., 2009) and the sensitivity (Garedew et al., 2004). Growth in the calorimetric set up was verified to be equivalent to that of a control cell culture (kept in an Erlenmeyer flask, under orbital stirring, inside an incubator, at 30 °C) until the end of the exponential phase, which was the period investigated in this work. After the exponential phase, no diauxic shift was observed for cultures grown in the calorimetric setting, probably due to oxygenation problems in the circuit that pumped cells through the calorimeter.

The flow line and the calorimetric vessel were sterilized after each experiment by circulating: *(i)* a 20 % (v/v) bleach aqueous solution, for at least, 4-5 h; *(ii)* deionized water for 10-20 min; (*iii*) 70 % (v/v) aqueous ethanol solution for 20 min; and *(iv)*deionized water for 10-20 min. Alternatively the system was sterilized by circulating a 70 % (v/v) aqueous ethanol solution for at least 3 h and then circulating H_2_O for 1 h to wash cellular debris. The reaction vessel and other glassware were sterilized at 170 °C during 2.5 h.

Electrical calibrations of the calorimetric system under the same conditions used in the main experiments (e.g. the pump flow rate, temperature) were performed as previously described (Leskiv et al., 2009). A value of *ε* = 15.5±0.2 μW·μV^-1^ (*n* = 41) for the electric calibration was used throughout this work.

In order to assign the power dissipated by a cell culture to a specific cell number or biomass it is necessary to know the effective volume of the calorimetric cell (O’Neill et al., 2003), which differs from its physical volume since the flow of cell culture results in transport of heat away from the detection area. The effective volume of the LKB 10700-1 flow-through cell, *V_eff_* = 0.547±0.012 mL (*n* = 4), under the temperature and volume flow rate conditions of the present experiments, was determined as recommended in the literature (O’Neill et al., 2003), using the kinetics of the base catalyzed hydrolysis of methyl paraben (Bento et al., 2018). The calculations also relied on the standard molar enthalpy of that reaction at 30 °C, 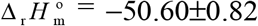 kJ·mol^-1^, proposed as benchmark (Bento et al., 2016). Experiment/calibration programing and data acquisition, were performed with the CBCAL software (Bernardes, 2022).

To obtain the specific ED rate, corresponding to power divided by the number of cells, the number of *S. cerevisiae* cells flowing through the calorimetric sensor at each time point is needed. The logarithmic of OD_600_ data was fitted to a polynomial equation, which was then used to obtain *OD*_600_ values for the culture used in the calorimetric measurements, during growth. Alternatively, the *OD*_600_ was followed in real-time by using an in-house built light dispersion system (LDS), composed by a white LED, that introduces light into the cell culture vessel, and a light controlled resistor (LDR GL5528), which measures the light intensity decay due to the increase of the number of cells along the growth process. The optical path of the system was 30 mm, and in this case an exponential smoothing was applied to the *OD*_600_ results to reduce the noise of the sensor readings before. Results obtained from both procedures were similar, namely, 1.4% and 6.5% deviations for SC and YPD media, respectively.

### MS-based untargeted metabolomics

An untargeted metabolomics approach was employed to compare the metabolomic brain profile of test samples with that of control samples. HRMS spectra were acquired on an Impact II QqTOF MS (Bruker Daltoniks) equipped with an electrospray source, interfaced in-line with an Elute UHPLC (Bruker Daltoniks). For each sample, 3 full scans and 1 auto MS/MS scan were performed using both positive and negative mode ionisation.

Reverse-phase chromatographic separations were performed on a Luna 2.5 μm C18(2)-HST column (100Å, 150 × 2 mm, Phenomenex) at a constant temperature of 40□°C, using a gradient elution of 0.1% formic acid in water (mobile phase A) and 0.1% formic acid in acetonitrile (mobile phase B) at a flow rate of 250□μL/min. The gradient conditions were as follows: 0.0-0.5 min 0 % B, 0.5-1.5 min 0 to 20 % B, 1.5-4.0 min 20 to 60 % B, 4.0-6.0 min 60 to 100 % B, 6.0-9.0 min 100 % B, 9.0-10.0 min 100 to 0 % B, 10.0-15.0 min 0 % B. For hydrophilic interaction chromatography (HILIC), a XBridge BEH Amide XP Column, (130Å, 2.5 μm, 150 x 2.1 mm, Waters) was used at a constant temperature of 40 °C, with a gradient elution of 10mM ammonium acetate in water containing 0.1% acetic acid (A) and 10mM ammonium acetate in acetonitrile containing 2% of water and 0.1% acetic acid (B) at a flow rate of 250 μL/min. The gradient conditions were as follows: 0-2 min 90 % B, 2-6 min 90 to 70 % B, 6-9 min 70 to 30 % B, 9-13 min 30 % B, 13-18 min 30 to 90 % B, 18-22 min 90 % B.

Mass spectrometer parameters were for full scan analysis were as follows: capillary voltage 3 kV (ESI+) or 4.5 kV (ESI-), end plate offset 500 V, nebulizer 4.0 bar, dry gas flow 8.0 L/min, dry heater temperature 200 °C. Spectra acquisition was performed with an absolute threshold of 25 counts per 1000. For auto MS/MS data acquisition, capillary voltage was set at 4.5 kV (ESI+) or 3.5 kV (ESI-), with an end plate offset of 500 V, nebulizer pressure of 4.0 bat, dry gas flow of 8.0 L/min and a heater temperature of 200 °C. Spectra acquisition was performed with a threshold of 20 counts per 1000, a cycle time of 3.0 seconds with exclusion after 3 spectra and release after 1.00 min. Internal calibration was achieved with a sodium formate/acetate solution introduced to the ion source ***via*** a 20 μL loop at the beginning of each analysis. Calibration was then performed using high-precision calibration mode (HPC). All acquisitions were performed with a *m/z* range from 50 to 1300 a.m.u.

The acquired MS data were processed by Data Analysis 4.1 (Bruker Daltoniks). For the identification of *in vivo* metabolites in the metabolomics approach, raw data were converted to mzXML format using ProteoWizard MSConvert (Chambers et al., 2012). Metabolite identification and statistical analysis was performed using XCMS (Benton et al., 2010; Benton et al., 2008; Forsberg et al., 2018; Smith et al., 2006) and the set of parameters given in Table 1 of SI, which were adapted from the original XCMS parameters for Bruker’s Q-TOF spectrometers. Metabolites identified by METLIN (Smith et al., 2005) were grouped into pathways in order to identify up- and down-regulated metabolites between samples from SC and eSC-derived samples. Metabolite abundance comparison between samples/conditions was performed using the heteroscedastic Welch *t*-test.

## 1 Supplementary Information

**Table 1.**
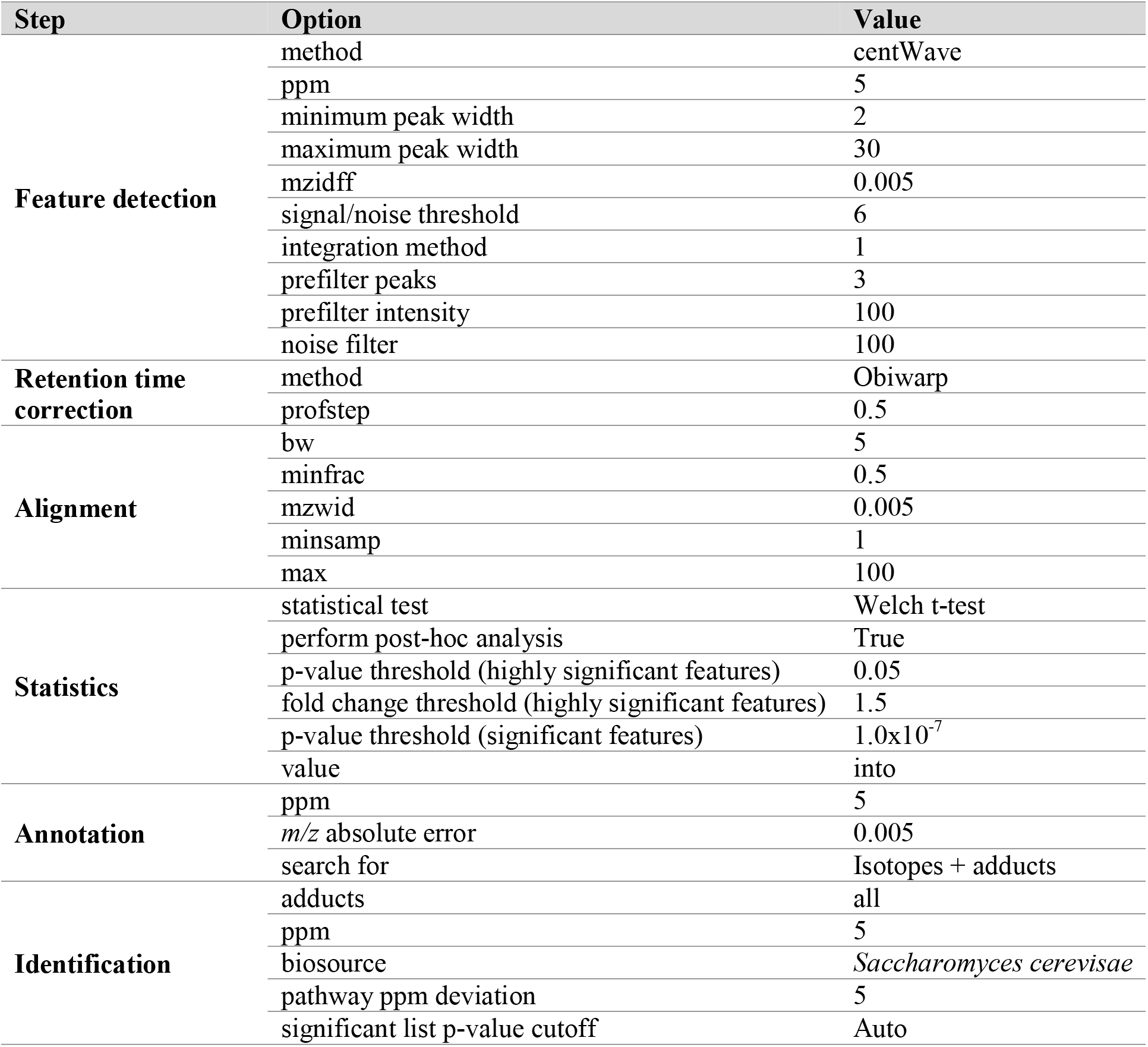
Parameters used in XCMS v3.7.1 analysis for yeast untargeted metabolomics data process.

**Figure SI1.**
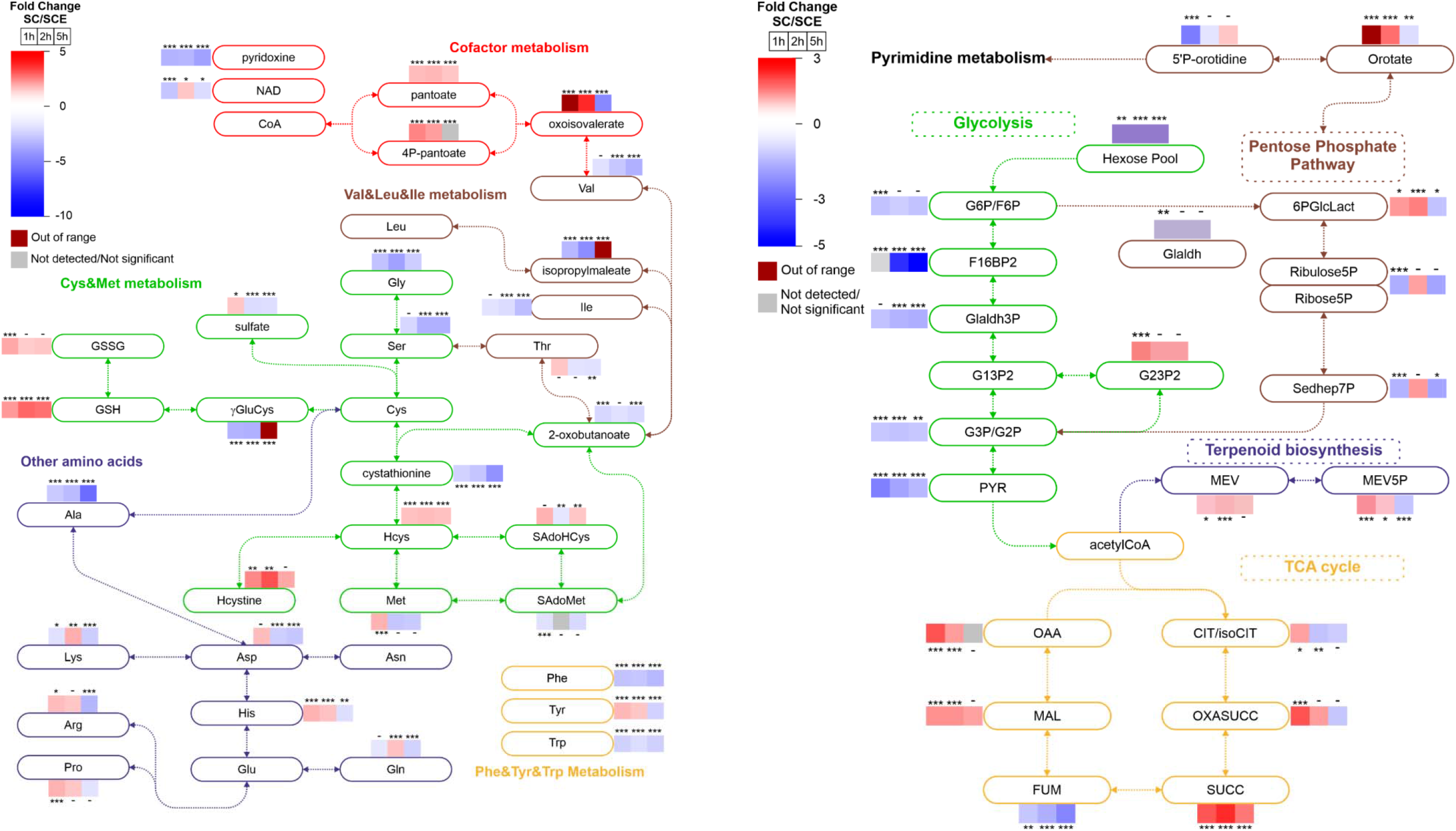
Simplified metabolic maps showing Log2 fold-changes of intracellular metabolite levels for cultures grown in Sc compared with eSC media during the lag phase (1 h), at the transition from the lag to the exponential phase (2 h), and at the exponential phase (5 h).). ***, p < 0.02; **, p < 0.05; *, p < 0.1. 6PGlcLact, 6P-gluconolactone; CIT, citrate; F16BP2, fructose-1,6-bisP; FUM, fumarate; G13P2, glyceG6P/F6P, glucose/fructose-6-P; G13P2/G23P2, glycerate-1,3/2,3-bisP; G3P/G2P, glycerate-3/2P;Gladh, glyceraldehyde; Gladh3P, glyceraldehyde-3P; MAL, malate; MEV, mevalonate; OAA, oxaloacetate; OXASUCC, oxalosuccinate; PYR, pyruvate; Sedhep7P, sedoheptulose-7P; SUCC, succinate; HCys, homocysteine; Hcystine, homocysteine; SAdoHCys, S-adenosyl-homocysteine; SAdoMet, S-adenosyl-methionine.

## Notes

### Competing Interest Statement

The authors have declared no competing interest.

## REFERENCES

Beard, D.A., and Qian, H. (2007). Relationship between Thermodynamic Driving Force and One-Way Fluxes in Reversible Processes. PLoS ONE 2, e144.

Beezer, A.E., ed. (1980). Biological Microcalorimetry. (London: Academic Press).

Bento, R.N., Rendas, M.A., Bernardes, C.E.S., Santos, M.S.C.S., Antunes, F., and Minas da Piedade, M.E. (2018). Kinetics of the base catalysed hydrolysis of methyl paraben revisited: Implications for determination of the effective volume of flow-microcalorimeters used to study cell cultures. Thermochim. Acta 659, 82–88.

Bento, R.N., Rendas, M.A., Semedo, V.A.R., Bernardes, C.E.S., Santos, M.S.C.S., Diogo, H.P., Antunes, F., and Minas da Piedade, M.E. (2016). The standard molar enthalpy of the base catalysed hydrolysis of methyl paraben revisited. J. Chem. Thermodyn. 103, 176–180.

Benton, H.P., Want, E.J., and Ebbels, T.M.D. (2010). Correction of mass calibration gaps in liquid chromatography-mass spectrometry metabolomics data. Bioinformatics 26, 2488–2489.

Benton, H.P., Wong, D.M., Trauger, S.A., and Siuzdak, G. (2008). XCMS2: Processing tandem mass spectrometry data for metabolite identification and structural characterization. Anal. Chem. 80, 6382–6389.

Bernardes, C.E.S. (2022). CBCAL: A Data Collection Program for Calorimetry Experiments (Version 3). Zenodo, https://doi.org/10.5281/zenodo.6475251.

Blomberg, A., Larsson, C., and Gustafsson, L. (1988). Microcalorimetric Monitoring of Growth of Saccharomyces-Cerevisiae - Osmotolerance in Relation to Physiological-State. J. Bacteriol. 170, 4562–4568.

Brachmann, C.B., Davies, A., Cost, G.J., Caputo, E., Li, J.C., Hieter, P., and Boeke, J.D. (1998). Designer deletion strains derived from Saccharomyces cerevisiae S288C: a useful set of strains and plasmids for PCR-mediated gene disruption and other applications. Yeast 14, 115–132.

Branco, M.R., Marinho, H.S., Cyrne, L., and Antunes, F. (2004). Decrease of H_2_O_2_ plasma membrane permeability during adaptation to H2O2 in Saccharomyces cerevisiae. J. Biol. Chem. 279, 6501–6506.

Brettel, R., Lamprecht, I., and Schaarschmidt, B. (1980). Micro-Calorimetric Investigations of the Metabolism of Yeasts.7. Flow-Calorimetry of Aerobic Batch Cultures. Radiat. Environ. Biophys. 18, 301–309.

Chambers, M.C., Maclean, B., Burke, R., Amodei, D., Ruderman, D.L., Neumann, S., Gatto, L., Fischer, B., Pratt, B., Egertson, J., et al. (2012). A cross-platform toolkit for mass spectrometry and proteomics. Nat. Biotechnol. 30, 918–920.

Cohen, R., and Engelberg, D. (2007). Commonly used Saccharomyces cerevisiae strains (e.g. BY4741, W303) are growth sensitive on synthetic complete medium due to poor leucine uptake. FEMS Microbiol. Lett. 273, 239–243.

Corbacho, I., Teixido, F., Velazquez, R., Hernandez, L.M., and Olivero, I. (2011). Standard YPD, even supplemented with extra nutrients, does not always compensate growth defects of Saccharomyces cerevisiae auxotrophic strains. Anton Leeuw Int. J. Gen. Mol. Biol. 99, 591–600.

DeBerardinis, R.J., and Thompson, C.B. (2012). Cellular Metabolism and Disease: What Do Metabolic Outliers Teach Us? Cell 148, 1132–1144.

Forsberg, E.M., Huan, T., Rinehart, D., Benton, H.P., Warth, B., Hilmers, B., and Siuzdak, G. (2018). Data processing, multi-omic pathway mapping, and metabolite activity analysis using XCMS Online. Nat. Protoc. 13, 633–651.

Garedew, A., Schmolz, E., and Lamprecht, I. (2004). Microcalorimetric investigation on the antimicrobial activity of honey of the stingless bee Trigona spp. and comparison of some parameters with those obtained with standard methods. Thermochim. Acta 415, 99–106.

Hanscho, M., Ruckerbauer, D.E., Chauhan, N., Hofbauer, H.F., Krahulec, S., Nidetzky, B., Kohlwein, S.D., Zanghellini, J., and Natter, K. (2012). Nutritional requirements of the BY series of Saccharomyces cerevisiae strains for optimum growth. FEMS Yeast Res. 12, 796–808.

Harrison, R., Papp, B., Pal, C., Oliver, S.G., and Delneri, D. (2007). Plasticity of genetic interactions in metabolic networks of yeast. Proc. Natl. Acad. Sci. USA 104, 2307–2312.

Hartman, J.L., Garvik, B., and Hartwell, L. (2001a). Cell biology - Principles for the buffering of genetic variation. Science 291, 1001–1004.

Hartman, J.L.t., Garvik, B., and Hartwell, L. (2001b). Principles for the buffering of genetic variation. Science 291, 1001–1004.

James, A.M., ed. (1987). Thermal and Energetic Studies of Cellular Biological Systems. (Bristol: IOP Publishing).

Jost, M., Santos, D.A., Saunders, R.A., Horlbeck, M.A., Hawkins, J.S., Scaria, S.M., Norman, T.M., Hussmann, J.A., Liem, C.R., Gross, C.A., et al. (2020). Titrating gene expression using libraries of systematically attenuated CRISPR guide RNAs. Nature Biotechnology 38, 355–+.

Keren, L., Hausser, J., Lotan-Pompan, M., Slutskin, I.V., Alisar, H., Kaminski, S., Weinberger, A., Alon, U., Milo, R., and Segal, E. (2016). Massively Parallel Interrogation of the Effects of Gene Expression Levels on Fitness. Cell 166, 1282–+.

Lan, G., Sartori, P., Neumann, S., Sourjik, V., and Tu, Y.H. (2012). The energy-speed-accuracy trade-off in sensory adaptation. Nat. Phys. 8, 422–428.

Leskiv, M., Bernardes, C.E.S., and Minas da Piedade, M.E. (2009). A calorimetric system based on the LKB 10700-1 flow microcalorimeter. Meas. Sci. Technol. 20, Artn 075107.

Melnykov, A.V. (2016). New mechanisms that regulate Saccharomyces cerevisiae short peptide transporter achieve balanced intracellular amino acid concentrations. Yeast 33, 21–31.

O’Neill, M.A.A., Beezer, A.E., Labetoulle, C., Nicolaides, L., Mitchell, J.C., Orchard, J.A., Connor, J.A., Kemp, R.B., and Olomolaiye, D. (2003). The base catalysed hydrolysis of methyl paraben: a test reaction for flow microcalorimeters used for determination of both kinetic and thermodynamic parameters. Thermochim. Acta 399, 63–71.

Orr, B.O., Hauswirth, A.G., Celona, B., Fetter, R.D., Zunino, G., Kvon, E.Z., Zhu, Y.W., Pennacchio, L.A., Black, B.L., and Davis, G.W. (2020). Presynaptic Homeostasis Opposes Disease Progression in Mouse Models of ALS-Like Degeneration: Evidence for Homeostatic Neuroprotection. Neuron 107, 95–+.

Papp, B. (2007). Plasticity of genetic interactions in metabolic networks of yeast. Febs J. 274, 244–244.

Proust, L., Sourabie, A., Pedersen, M., Besancon, I., Haudebourg, E., Monnet, V., and Juillard, V. (2019). Insights Into the Complexity of Yeast Extract Peptides and Their Utilization by Streptococcus thermophilus. Front. Microbiol. 10.

Robador, A., LaRowe, D.E., Finkel, S.E., Amend, J.P., and Nealson, K.H. (2018). Changes in Microbial Energy Metabolism Measured by Nanocalorimetry during Growth Phase Transitions. Front. Microbiol. 9, Article 109.

Schaarschmidt, B., Zotin, A.I., Brettel, R., and Lamprecht, I. (1975). Experimental Investigation of Bound Dissipation Function - Change of Psi-U-Function during Growth of Yeast. Archives of Microbiology 105, 13–16.

Schiraldi, A. (1995). Microbial-Growth and Metabolism - Modeling and Calorimetric Characterization. Pure Appl. Chem. 67, 1873–1878.

Smith, C.A., O’Maille, G., Want, E.J., Qin, C., Trauger, S.A., Brandon, T.R., Custodio, D.E., Abagyan, R., and Siuzdak, G. (2005). METLIN: a metabolite mass spectral database. Ther. Drug Monit. 27, 747–751.

Smith, C.A., Want, E.J., O’Maille, G., Abagyan, R., and Siuzdak, G. (2006). XCMS: Processing mass spectrometry data for metabolite profiling using Nonlinear peak alignment, matching, and identification. Anal. Chem. 78, 779–787.

Ulitsky, I., and Shamir, R. (2007). Pathway redundancy and protein essentiality revealed in the Saccharomyces cerevisiae interaction networks. Mol. Syst. Biol. 3.

Wishart, D.S., Bartok, B., Oler, E., Liang, K.Y.H., Budinski, Z., Berjanskii, M., Guo, A.C., Cao, X., and Wilson, M. (2021). MarkerDB: an online database of molecular biomarkers. Nucleic Acids Res. 49, D1259–D1267.

Woronoff, G., Nghe, P., Baudry, J., Boitard, L., Braun, E., Griffiths, A.D., and Bibette, J. (2020). Metabolic Cost of Rapid Adaptation of Single Yeast Cells Under Unforeseen Challenge. Proc. Natl. Acad. Sci. USA 117, 10660–10666.

