## Supplemental Information for "Energy dissipation in early detection of cellular responses to metabolic challenges"

### Supplementary Information

**Table 1.** Parameters used in XCMS v3.7.1 analysis for yeast untargeted metabolomics data process.

| Step | Option | Value |
| --- | --- | --- |
| Feature detection | method | centWave |
|  | ppm | 5 |
|  | minimum peak width | 2 |
|  | maximum peak width | 30 |
|  | mzidff | 0.005 |
|  | signal/noise threshold | 6 |
|  | integration method | 1 |
|  | prefilter peaks | 3 |
|  | prefilter intensity | 100 |
|  | noise filter | 100 |
| Retention time correction | method | Obiwarp |
|  | profstep | 0.5 |
| Alignment | bw | 5 |
|  | minfrac | 0.5 |
|  | mzwid | 0.005 |
|  | minsamp | 1 |
|  | max | 100 |
| Statistics | statistical test | Welch t-test |
|  | perform post-hoc analysis | True |
|  | p-value threshold (highly significant features) | 0.05 |
|  | fold change threshold (highly significant features) | 1.5 |
|  | p-value threshold (significant features) | 1.0x10^-7^ |
|  | value | into |
| Annotation | ppm | 5 |
|  | *m/z* absolute error | 0.005 |
|  | search for | Isotopes + adducts |
| Identification | adducts | all |
|  | ppm | 5 |
|  | biosource | *Saccharomyces cerevisae* |
|  | pathway ppm deviation | 5 |
|  | significant list p-value cutoff | Auto |

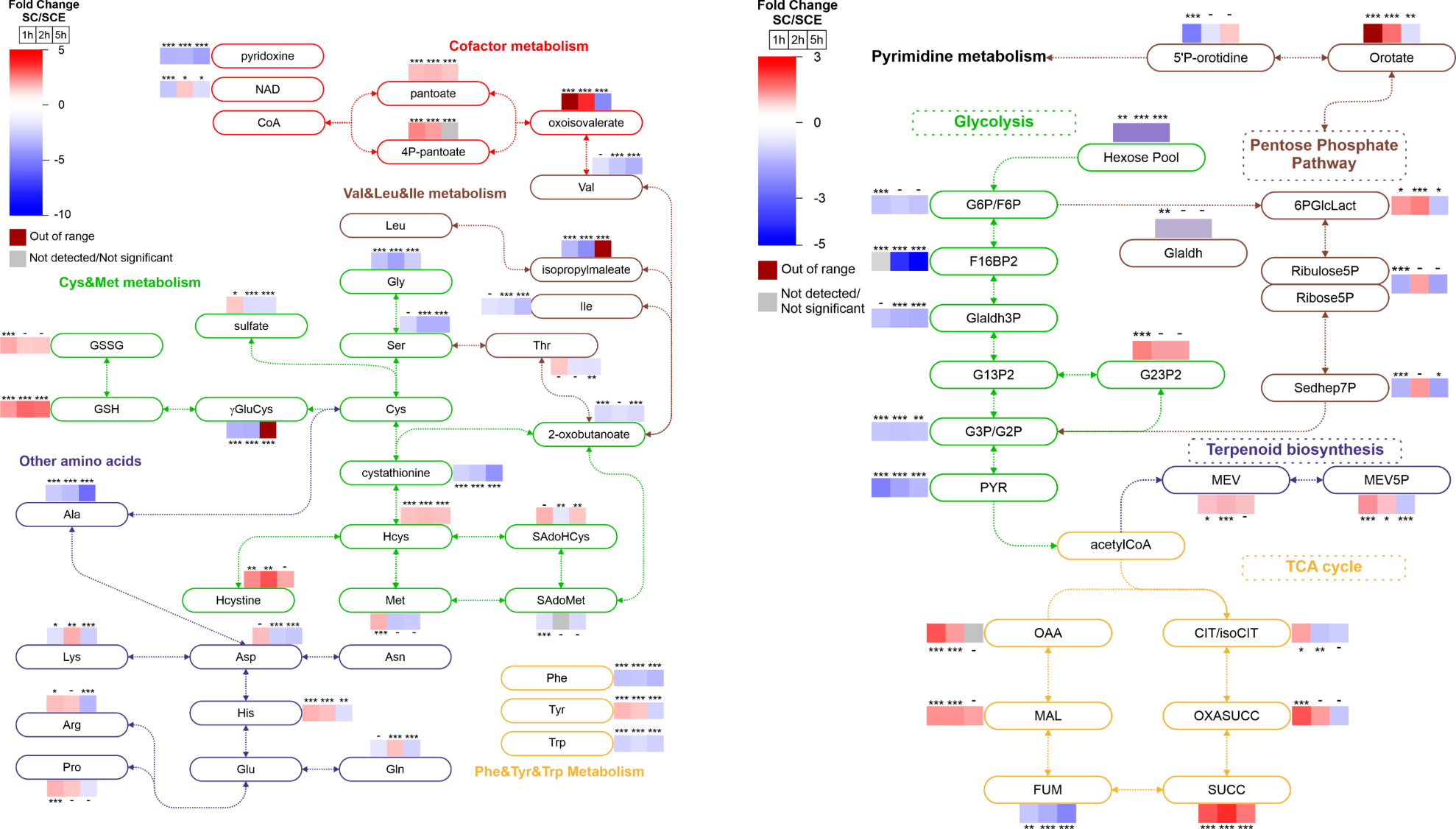

Figure SI1 – Simplified metabolic maps showing Log2 fold-changes of intracellular metabolite levels for cultures grown in Sc compared with eSC media during the lag phase (1 h), at the transition from the lag to the exponential phase (2 h), and at the exponential phase (5 h). ). ***, p < 0.02; **, p < 0.05; *, p < 0.1. 6PGlcLact , 6P-gluconolactone; CIT, citrate; F16BP2, fructose-1,6-bisP; FUM, fumarate; G13P2, glyceG6P/F6P, glucose/fructose-6-P; G13P2/G23P2, glycerate-1,3/2,3-bisP; G3P/G2P, glycerate-3/2P; Gladh, glyceraldehyde; Gladh3P, glyceraldehyde-3P; MAL, malate; MEV, mevalonate; OAA, oxaloacetate; OXASUCC, oxalosuccinate; PYR, pyruvate; Sedhep7P, sedoheptulose-7P; SUCC, succinate; HCys, homocysteine; Hcystine, homocysteine; SAdoHCys, S-adenosyl-homocysteine; SAdoMet, S-adenosyl-methionine.
